# Key Promoter Region of *Wnt4* response to FSH and Genetic Effect on Several Production Traits of Its Mutations in Chicken

**DOI:** 10.1101/2021.05.11.443679

**Authors:** Conghao Zhong, Yiya Wang, Cuiping Liu, Yunliang Jiang, Li Kang

**Author notes:** Correspondence, **Corresponding author:** Li Kang, Ph. D and Professor, Contacting information: College of Animal Science and Veterinary Medicine, Shandong Agricultural University, No. 61 Daizong street, Taian 271018, China. These authors have contributed equally to this work (Conghao Zhong, Yiya Wang).

## Abstract

The signaling pathway of the wingless-type mouse mammary tumor virus integration site (Wnt) plays an important role in ovarian and follicular development. *Wnt4* was shown in our previous study to be involved in the selection and development of chicken follicles by up-regulating the expression of follicle stimulating hormone receptors (*FSHR*), stimulating the proliferation of follicular granulosa cells and increasing the secretion of steroidal hormones. To further characterize cis-elements regulating chicken *Wnt4* transcription, in this study we determined critical regulatory regions affecting chicken *Wnt4* transcription, then identified a single nucleotide polymorphism (SNP) in this region, and finally analyzed the association of the SNP with chicken production traits. The results showed that the 5′ regulatory region from -3354 to -2689 of the chicken *Wnt4* gene had the strongest activity and greatest response to FSH stimulation, and that one SNP site -3015 (G > C) in this segment was identified as affecting the binding of NFAT5 (nuclear factor of activated T cells 5). When G was replaced by C at this site, it eliminated the binding by NFAT5. Moreover, this locus was significantly associated with the keel length and comb length of hens. Individuals with the genotype CG had longer keels while those with genotype CC had longer combs. Collectively, these data suggested that the SNP-3015 (G > C) is (i) involved in the regulation of *Wnt4* gene expression by affecting the binding of NFAT5, (ii) associated with chicken keel length and comb length, and (iii) is a potential DNA marker in the molecular breeding of chickens for egg laying.

## 1. INTRODUCTION

As a conservative signaling pathway, Wnt signaling regulates multiple developmental processes and occurrences of disease, such as stem cell self-renewal, cell proliferation, cell fate determination and early embryonic development and differentiation (Waghmare and Page-McCaw, 2018; Komiya and Habas, 2008; Cadigan and Nusse, 1997). The Wnt family members are a class of secreted glycoprotein signaling molecules with localized action, which are generally used as ligands to participate in signal transduction. There are 15 or even more receptors or co-receptors, such as the Frizzled (Fzd) protein family, that are recognized and bound by Wnt ligands (van Amerongen and Nusse, 2009; Niehrs, 2012). Two signaling pathways - β-catenin-dependent (canonical) and β-catenin-independent (non-canonical) - are used by Wnt ligands (Schwarz-Romond, 2012, van Amerongen et al., 2008).

Some Wnts and their homologous receptor components are expressed in postnatal ovaries but their role in ovarian physiology is still unclear. *Wnt4* plays an important regulatory function in adult ovarian follicles. Overexpression of *Wnt4* in granulosa cells of eCG-treated mice up-regulates the expression of β-catenin and key genes *CYP11A1, CYP19A1* and *StAR* in the synthesis of gonadal steroid hormones (Boyer et al., 2010). In chicken, *Wnt4* affects the growth, differentiation and development of the oviduct (Lim et al., 2013), and is mainly expressed in the shell glands and isthmus of the chicken oviduct, regulated by estrogen (Dougherty and Sanders, 2005). Our previous study revealed that the expression of *Wnt4* was the highest in the granulosa cells of small yellow follicles in chickens and declined in hierarchal follicles, and that *Wnt4* up-regulates the expression of *FSHR* and down-regulates the expression of *AMH* and *OCLN*, promotes the expression of *StAR* and *CYP11A1*, and stimulates the proliferation of granulosa cells (Wang et al., 2017).

In the current study, the regulatory mechanism of chicken *Wnt4* transcription was further investigated. The critical regulatory cis-elements responsible for *Wnt4* transcription that are also responsive to FSH treatment were first determined. Two single nucleotide polymorphisms (SNPs) in the 5′ regulatory region of the chicken *Wnt4* gene were identified, and their associations with production traits in hens were analyzed. Finally, the mechanism of the SNP that is associated with keel length and comb length was analyzed.

## 2. MATERIALS AND METHODS

### 2.1 Animals and Sample Collections

Three breeds of Hy-line brown hens, Jining Bairi hens and Sunzhi hens with different production performances were used in this study. Hens were randomly selected from the local farm affiliated with Shandong Agricultural University. The egg laying traits of the Jining Bairi population were recorded individually for association analysis. All chickens had free access to water and feed. The chickens were housed in separate cages with a daily light period of 16 h, and egg laying was monitored to determine the timing and regularity of laying. Genomic DNA was extracted from blood samples collected from the wing vein using a DNA extraction mini kit (Tiangen Biotech, Beijing, China). All sampled hens were killed by cervical dislocation immediately after oviposition and the abdominal cavity was opened. Preovulatory follicles were carefully collected from laying hens and placed in phosphate-buffered saline (PBS) with 1% penicillin/streptomycin for cell culture. All of the animal experiments were approved by the Institutional Animal Care and Use Ethics Committee of Shandong Agricultural University and performed in accordance with the “Guidelines for Experimental Animals” of the Ministry of Science and Technology of China.

### 2.2 Cell Culture

The hierarchical follicles were isolated from egg-laying hens and placed in PBS. The yolks of the follicles were removed carefully with ophthalmic forceps. The granulosa cells (GCs) were isolated from the hierarchical follicles and then dispersed by treatment with 0.25% trypsin-EDTA (Gibco, Camarillo, CA, USA) at 37 °C for 10 min with gentle oscillation in a centrifuge tube. After centrifugation, the GCs were suspended in a culture medium (M199 with 10% fetal bovine serum and 1% penicillin/streptomycin), and subsequently seeded in 24-well culture plates at a density of 1×10 ^5^/well. The number of cells was detected using Trypan blue. Cells were cultured at 38 °C in an atmosphere of water-saturated 5% CO_2_ for 24 h.

### 2.3 Construction of *Wnt4* Promoter Deletion Vectors and Site-directed mutation

The region from -3354 to +252 bp in the 5′-regulatory region of the chicken *Wnt4* promoter and five promoter deletion fragments, -2689/+252, -1875/+252, -1188/+252, -535/+252, were amplified from hen genomic DNA, where +1 is the transcription initiation site. Five forward primers contain the KpnI site at the ends, and one reverse primer located downstream to the transcription start site contains the HindIII site at the ends (primer sequences are listed in Table 1). All PCR fragments were digested with KpnI and HindIII restriction enzymes and ligated with pGL3-Basic vector (Promega, Madison, WI, USA).

**TABLE 1.**
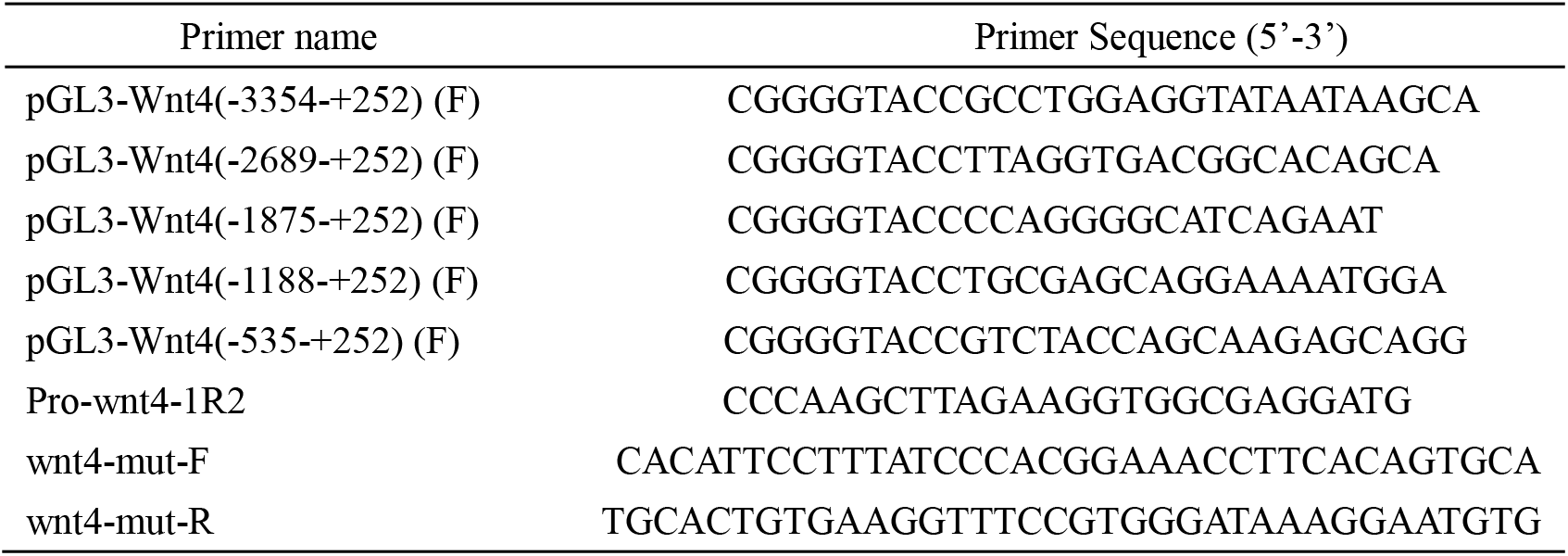
Primers used for plasmid construction of chicken *Wnt4* gene

Two plasmids including the wild type (pGL3-G) and mutation type (pGL3-C) were constructed to assess the functionality of this transcription factor binding site in the *Wnt4* promoter. The primers for g.-3015(G > C) mutations were designed using the -3354 to -2689 region of the *Wnt4* gene promoter as the template (primer sequences are listed in Table 1). The PCR products were digested by Dpn1 Methylase, so the plasmid template to be mutated could be removed and the plasmid containing the mutation site could be retained.

### 2.4 Cell Transfection and Luciferase Assay

GCs were plated on 24-well plates for transient transfection experiments using Lipofectamine LTX and Plus Reagent (Invitrogen). The cells were transfected with pGL4.74 control vector (Promega, Madison, WI, USA), and the five *Wnt4* luciferase plasmids differing in length (800 ng/well) along with recombinant FSH were added to the wells 6 h after transfection. In another experiment, the pGL4.74 control vector (Promega, Madison, WI, USA), the wild-type plasmid and the mutation-type plasmid (800 ng/well) were used to transfect cultured GC cells. Twenty-four hours after transfection, these cells were lysed for a luciferase activity assay.

Luciferase activity was measured using the Dual-Luciferase Reporter Assay System according to the manufacturer (Promega, Madison, WI, USA). The enzymatic activity of luciferase was measured with a luminometer (Modulus TM, Turner Biosystems). The individual values were averaged for each experiment, and the transfections were performed at least in triplicate. Empty pGL3-basic was used as the control. Luciferase activity was calculated by dividing the Firefly luciferase activity by the Renilla luciferase activity.

### 2.5 SNP Identification, Polymorphism and Association Analysis

Fifty individuals from Jining Bairi hens, Hy-line Brown hens, and Sunzhi hens were used as template for PCR amplification to the critical promoter region of *Wnt4* (−3353 to -2689), and then the amplifications were sequenced. The data, sequenced bidirectionally, were analyzed using the DNAMAN program (version 7.212, Lynnon Corp., Quebec, Canada) to determine the potential SNPs within these amplifications. Primer pairs amplified the -3353 to -2689 fragments (the primers are shown in Table 1). The genotypes at the -3015 SNP site were determined by Kompetitive Allele Specific PCR (KASP) (Baygene Biotechnology Co., Shanghai, China) in the Jining Bairi population. The genotype and allele frequencies and Hardy-Weinberg equilibrium P-value were calculated using the Tools for Population Genetic Analyses software (http://www.marksgeneticsoftware.net/tfpga.htm). The association of SNP with egg laying traits in the Jining Bairi population was analyzed using the following general model in SPSS (SPSS Inc., Chicago, IL, USA): Y_ij_ =*μ*+ G_i_ + e_ij_, where Y_ij_ is the phenotypic value of traits, *μ* is the population mean, G_i_ is the fixed effect of genotype, and e_ij_ is the random error effect.

### 2.6 Electrophoretic Mobility Shift Assay (EMSA)

The Genomatix software (www.genomatix.de) was used to predict that the -3015 (G > C) may be the transcription factor binding sites of NFAT5, which regulates the transcription of Wnt4 by responding to FSH. HIH3T3 cells were seeded at a density of 1 × 10^6^ cells/mL and incubated in DMEM with 10% FBS at 37 °C for 72 h. The nuclear extracts prepared from the cells were incubated with biotin-labeled double-stranded oligonucleotides containing the consensus sequences for NFAT5 (5′ -TTTATCCCAgGGAAACCTTCACAGTGCATTC-3′) for an additional 4 h. GATA1 (5′ CACTTGATAACAGAAAGTGATAACTCT-3′) was used as a control. EMSA was performed using Non-Radioactive EMSA Kits with Biotin-Probes (Viagene, Tampa, FL, USA). The DNA-protein complex and unbound probe were electrophoresed on a 6% native polyacrylamide gel and visualized as per western blotting. The NFAT5 monoclonal antibody was used for the super shift, and standard NFAT5 was used as a positive control.

### 2.7 Statistical Analyses

The experiments were repeated a minimum of three times using tissues from different hens. All data are presented as the means ± SEM. The differences between different groups were determined by one-way ANOVA followed by Duncan’s test in SPSS (SPSS Inc., Chicago, IL, USA). The differences between groups were considered statistically significant when *P* < 0.05.

## 3. RESULTS

### 3.1 Critical Region of *Wnt4* Gene response to FSH in Chicken GC Cells

As the *Wnt4* gene is reported to play an essential role in follicle selection (Wang et al., 2017), we set out to analyze the mechanisms regulating its transcription. The luciferase activity assay on chicken preovulatory follicle GC cells transfected with different *Wnt4* promoter vectors (Table 1) showed that deletion from -3354 to -2689 greatly decreased relative luciferase activity, indicating positive regulatory elements exist in this region and that the region from -3354 to -2689 had the greatest response to FSH (10 ng/mL) stimulation (Figure 1), suggesting cis-acting response elements to FSH exist to regulate chicken *Wnt4* transcription.

**FIGURE 1.**
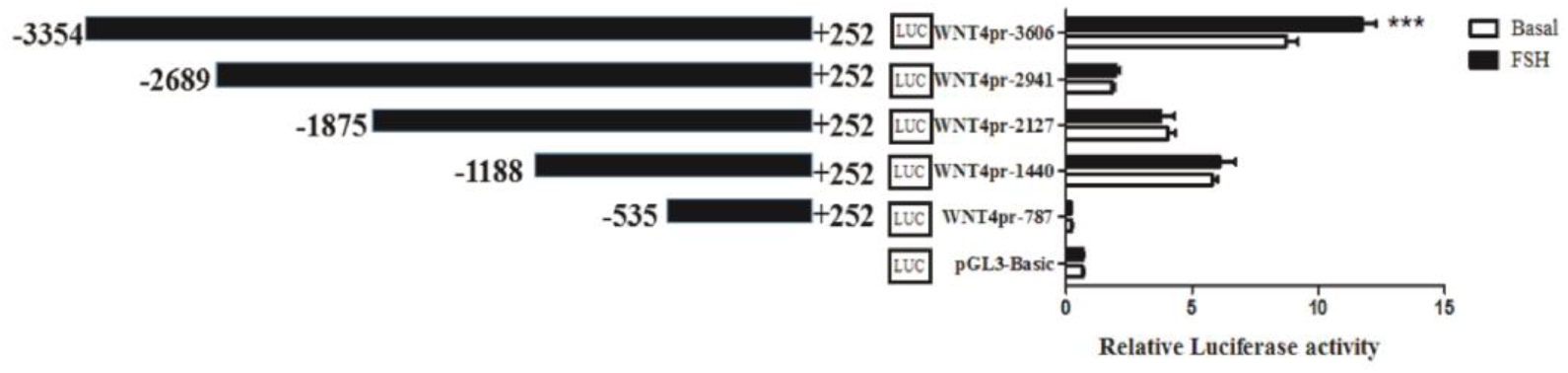
Luciferase assay of effecting of FSH on *Wnt4* promoter activity in chicken GCs. The numbering refers to the transcription initiation site designated as +1. Each bar represents the means ± SEM. *** represent *P* ≤ 0.001

### 3.2 Polymorphisms in the Critical Promoter Region of the Chicken *Wnt4* Gene

Sequence alignment between the promoter regions of the chicken *Wnt4* gene from Hy-line Brown, Jining Bairi and Sunzhi hens showed that the critical promoter region contains a SNP (G > C) at -3015 (Figure 2A). The peak map of polymorphic sites was genotyped using Chromas software. This polymorphic site has three genotypes (Figure 2B), and the number distribution of different genotypes is shown in Figure 2C. According to the results of KASP, the genotype frequency and allele frequency of this SNP locus were calculated (Table 2). At this SNP site, allele G was predominant in the Jining Bairi chicken population.

**FIGURE 2.**
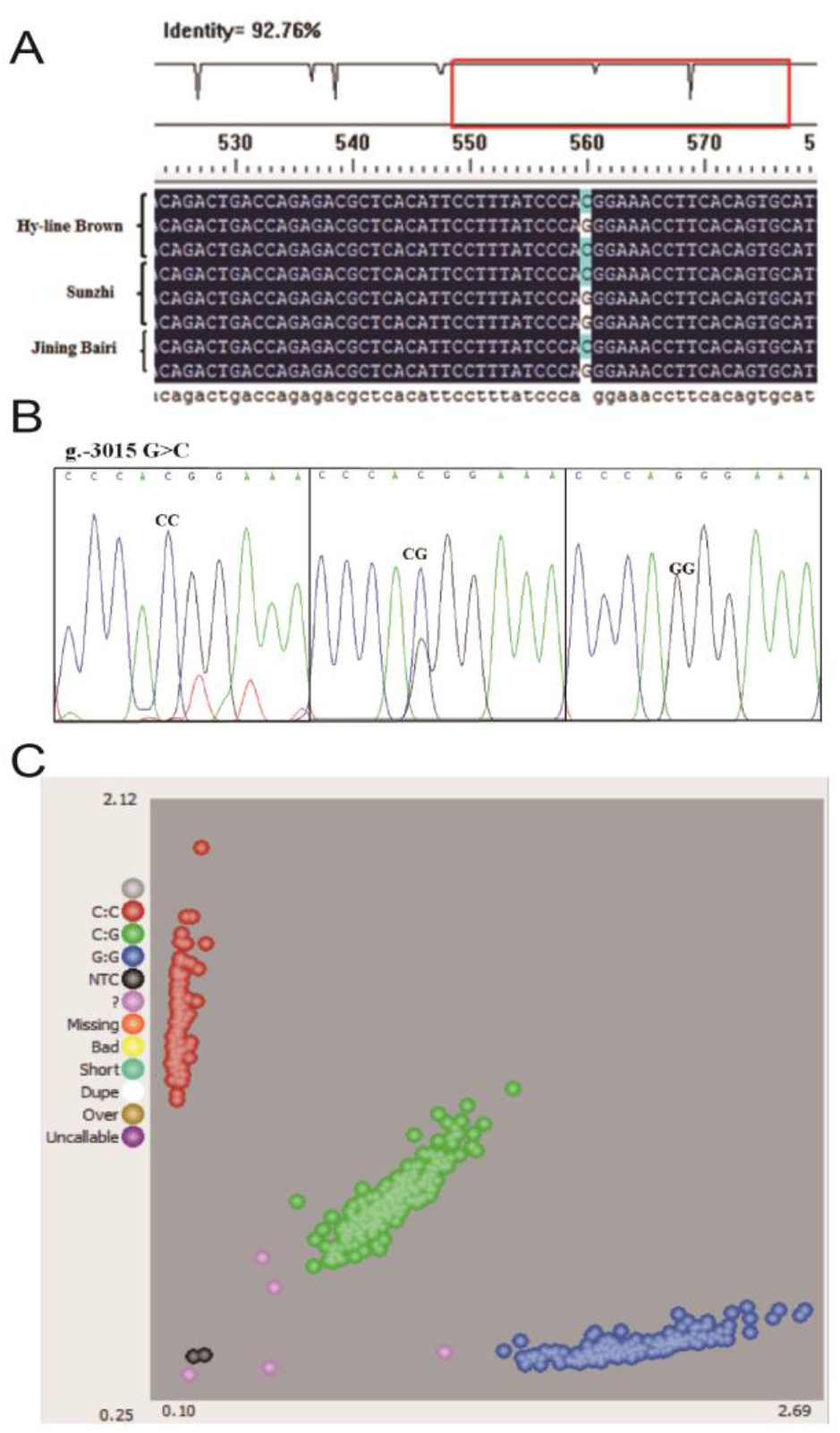
Polymorphisms in the critical promoter region of chicken *Wnt4* gene. (A) Sequencing alignment of SNP site in Hy-line Brown hens, Jining Bairi hens and Sunzhi hens and one SNP (g.-3015) was detected. (B) Genotyping of the SNP (g.-3015) site and three genotypes were showed. (C) Genotyping in Jining Bairi population (n = 539) by using the KASP method.

**TABLE 2.** Genotype and allele frequencies at the SNP site of the *Wnt4* gene in the Jining Bairi chicken

| Site | Breed (number) | Genotype frequency |  |  | Allele frequency |  | HWE (*P*-value) |
| --- | --- | --- | --- | --- | --- | --- | --- |
| g.-3015 | Jining Bairi<br>(n=539) | GG | CG | CC | G | C | 0.204 |
|  |  | 0.358 | 0.458 | 0.183 | 0.587 | 0.413 |  |

### 3.3 Association of the SNP-3015(G > C)of Wnt4 Gene with Chicken Production Traits

The statistical analysis is based on the genotype results of the Jining Bairi chicken population (n = 539) with production records. The association between the genotype of each individual and the egg laying traits is shown in Table 3. The results indicate that chickens with genotype CG have a longer keel than chickens with the other genotypes (*P* < 0.05) and that the CC individuals have a longer comb (*P* < 0.05). There was no significant difference between genotypes for the other measured traits.

**TABLE 3.** Associations of genotypes of the SNP on laying traits in the Jining Bairi chicken

| Site | Traits | Genotype |  |  | *P*-value |
| --- | --- | --- | --- | --- | --- |
|  |  | GG | CG | CC |  |
| g.-3015 | AfE | 149.763±0.411 | 150.518±0.409 | 150.235±0.764 | 0.473 |
|  | KLfE | 10.693±0.056<sup>ab</sup> | 10.734±0.048<sup>a</sup> | 10.531±0.068<sup>b</sup> | 0.024 |
|  | BWfE | 1463.082±17.766 | 1476.105±10.399 | 1475.234±13.269 | 0.330 |
|  | E30 | 111.602±1.690 | 113.539±1.074 | 113.083±1.346 | 0.411 |
|  | E50 | 137.520±2.039 | 140.223±1.293 | 138.818±1.702 | 0.075 |
|  | CLfE | 4.005±0.052<sup>a</sup> | 4.080±0.048<sup>ab</sup> | 4.224±0.080<sup>b</sup> | 0.019 |

Note: AfE: age of first egg; KLfE: keel length at the age of first egg; BWfE: body weight at the age of first egg; E30: egg number at 30 weeks of age; E50: egg number at 50 weeks of age, CLfE: comb length at the age of first egg.

### 3.4 Effect of the SNP-3015(G > C)on Promoter Activity of the Chicken *Wnt4* Gene

Luciferase reporter constructs of pGL3-G and pGL3-C were transiently transfected into GCs to assess whether the SNP could change the effect of *Wnt4* gene transcription. As shown in Figure 3, this SNP significantly affected the promoter activity - the promoter with allele G has higher luciferase activity than the promoter with allele C (*P* ≤0.001).

**FIGURE 3.**
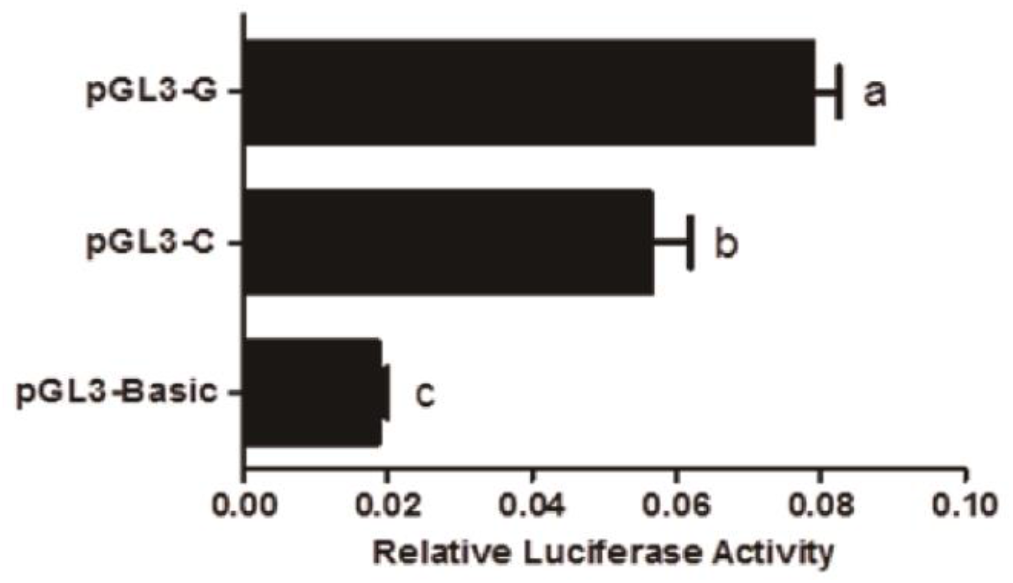
Effect of the SNP on chicken *Wnt4* gene transcriptional activity in GCs. It used the dual-luciferase report assay for verification of the effect of the SNP (g.-3015) different genotypes. Small letters on the error bar represent *P* ≤ 0.05

### 3.5 SNP-3015(G > C)Affects NFAT5 Binding in Chicken *Wnt4* Promoter

Analysis with the Genomatix revealed that the SNP (G > C) at -3015 may affect the binding site of NFAT5, which may be related to the regulation of *Wnt4* responses to FSH. An electrophoretic mobility shift assay (EMSA) was performed to identify this transcription factor. As shown in Figure 5, binding to the oligo nucleotide (NFAT5) was detected when the SNP site was G (Figure 4A). The super shift assay with the NFAT5 monoclonal antibody appeared to correspond to the DNA-protein-antibody complex, which further demonstrated that the SNP (g.-3015) was located in the binding site of NFAT5 (Figure 4B).

**FIGURE 4.**
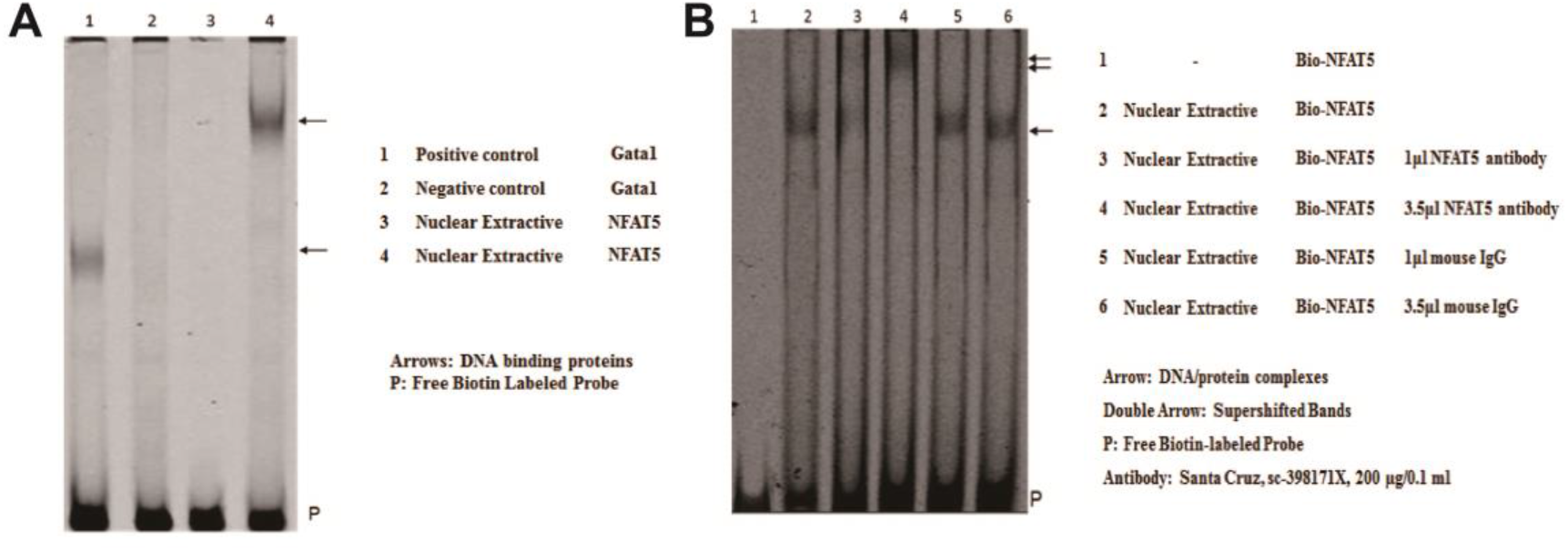
Electrophoretic mobility shift assay (EMSA) of the transcription factor binding sites at the SNP (g.-3015) in chicken *Wnt4* gene promoter. (A) EMSA analysis was conducted with biotin-labeled probe in the SNP (g.-3015). (B) Super shift assay with NFAT5 monoclonal antibody appeared to correspond to the DNA-protein-antibody complex.

## 4. DISCUSSION

Our previous work has proved that *Wnt4* plays an important role in ovarian follicle selection in chickens, and FSH treatment significantly increased the expression of *Wnt4* in GCs (Wang et al., 2017). In cows, the *Wnt4* signal can also enhance the stimulation of FSH during the follicular dominance phase (Gupta et al., 2014). These results suggest that there is interaction between *Wnt4* and FSH. However, the molecular mechanism of the interaction between them has not been investigated. In our study, we found that the 5′ regulatory region from -3354 to -2689 of *Wnt4* was the key regulatory region in response to FSH stimulation and affected chicken *Wnt4* gene transcription, which provides a reference for further elucidation of the regulatory relationship between *Wnt4* and FSH.

There are two studies showing that SNPs of the *Wnt4* gene can affect its expression and function. SNP rs7521902 in *Wnt4* is significantly correlated with the pathogenesis of endometriosis (Angioni et al., 2020), and rs2072920 in *Wnt4* is associated with body mass index (BMI) variations in the Han Chinese population (Dong et al., 2017). We identified the critical regions of the *Wnt4* promoter which contained a SNP -3015 (G > C) and we showed that SNP -3015 (G > C) in the *Wnt4* gene was significantly associated with keel length and comb length in chickens, the individuals with CG genotype having a longer keel length.

The Wnt pathway is involved in bone and cartilage homeostasis, controls the function of osteoblasts, and affects the differentiation of osteoblasts (Huang et al., 2019; Guo and Cooper, 2007; Li et al., 2019). A genome-wide association study revealed that polymorphisms of *Wnt4* were associated with bone mineral densities (Hendrickx et al., 2017), which is consistent with the result of our study. Keel bone health in laying hens is an important welfare problem in egg production systems. Keel bone damage (KBD) has adverse effects on welfare, health, production performance, and egg quality of the laying hens and is regulated by genetic factors (Eusemann et al., 2020). There is a strong association between egg production and keel bone fractures (KBF); the laying performance of KBF hens is 16.2% lower than that of non-fractured hens (Rufener, 2019). Although the relationship between keel length and KBF or KBD is still unclear, the length of keel affects the development of internal organs to a certain extent in domestic chickens. Therefore, selective breeding may help to reduce the susceptibility of KBF (Stratmann et al., 2016; Kolakshyapati et al., 2019). The SNP -3015 (G > C) in the chicken *Wnt4* promoter is a useful molecular marker for breeding focused on improving the keel trait. Comb size is used as an important criterion for hen development and egg production (Folsch et al., 1994; Emily et al., 2011). In our study, we found that individuals with the CC genotype had a longer comb at the -3015 (G > C) mutation site, suggesting a potential marker for improving the comb trait.

The transcription factor prediction and EMSA experiment showed that the SNP-3015 (G > C) in this key regulatory region was the binding site of the transcription factor NFAT5. When this site is G, it has binding activity. In mice, NFAT5 can regulate the proliferation of granulosa cells by activating the Wnt signaling pathway (Tao et al., 2019). We suppose that the SNP -3015 (G > C) may regulate *Wnt4* expression through NFAT5 binding, in turn further affecting chicken follicle selection; this possibility requires further investigation.

In conclusion, SNP-3015 (G > C) in the *Wnt4* promoter region was regulated by NFAT5. The SNP was significantly associated with chicken keel and comb length. These data provide a reference for further elucidation of the relationship between the appearance of phenotypes such as keel length and comb length and reproductive capacity in chickens.

## ACKNOWLEDGEMENTS

This study was funded by the National Natural Science Foundation of China (31672414, 31772588), Agricultural Breed Project of Shandong Province (2019LZGC019), and the Funds of Shandong “Double Tops” Program (SYL2017YSTD12).

## CONFLICT OF INTEREST

The authors declare no conflict of interests for this article.

## REFERENCES

Angioni, S., D’Alterio, M. N., Coiana, A., Anni, F., Gessa, S., & Deiana, D. (2020). Genetic Characterization of Endometriosis Patients: Review of the Literature and a Prospective Cohort Study on a Mediterranean Population. Int J Mol Sci, 21(5). doi:10.3390/ijms21051765

Boyer, A., Lapointe, E., Zheng, X., Cowan, R. G., Li, H., Quirk, S. M., … Boerboom, D. (2010). WNT4 is required for normal ovarian follicle development and female fertility. FASEB J, 24(8), 3010–3025. doi:10.1096/fj.09-145789

Cadigan, K. M., & Nusse, R. (1997). Wnt signaling: a common theme in animal development. Genes Dev, 11(24), 3286–3305. doi:10.1101/gad.11.24.3286

Dong, S. S., Hu, W. X., Yang, T. L., Chen, X. F., Yan, H., Chen, X. D., … Guo, Y. (2017). SNP-SNP interactions between WNT4 and WNT5A were associated with obesity related traits in Han Chinese Population. Sci Rep, 7, 43939. doi:10.1038/srep43939

Dougherty, D. C., & Sanders, M. M. (2005). Estrogen action: revitalization of the chick oviduct model. Trends Endocrinol Metab, 16(9), 414–419. doi:10.1016/j.tem.2005.09.001

Emily A. O’Connor, John E. Saunders, Hannah Grist, Morven A. McLeman & Siobhan M. (2011). The relationship between the comb and social behaviour in laying hens. Abeyesinghe, 135(4), 293–299. doi: 10.1016/j.applanim.2011.09.011

Eusemann, B. K., Patt, A., Schrader, L., Weigend, S., Thone-Reineke, C., & Petow, S. (2020). The Role of Egg Production in the Etiology of Keel Bone Damage in Laying Hens. Front Vet Sci, 7, 81. doi:10.3389/fvets.2020.00081

Folsch, D. W., Sulzer, B., Meier, T., & Huber, H. U. (1994). [The effect of the husbandry system on comb size and comb color in hens]. Tierarztl Prax, 22(1), 47–54. Retrieved from https://www.ncbi.nlm.nih.gov/pubmed/8165660

Guo, J., & Cooper, L. F. (2007). Influence of an LRP5 cytoplasmic SNP on Wnt signaling and osteoblastic differentiation. Bone, 40(1), 57–67. doi:10.1016/j.bone.2006.07.016

Gupta, P. S., Folger, J. K., Rajput, S. K., Lv, L., Yao, J., Ireland, J. J., & Smith, G. W. (2014). Regulation and regulatory role of WNT signaling in potentiating FSH action during bovine dominant follicle selection. PLoS One, 9(6), e100201. doi:10.1371/journal.pone.0100201

Hendrickx, G., Boudin, E., Steenackers, E., Nielsen, T. L., Andersen, M., Brixen, K., & Van Hul, W. (2017). Genetic Screening of WNT4 and WNT5B in Two Populations with Deviating Bone Mineral Densities. Calcif Tissue Int, 100(3), 244–249. doi:10.1007/s00223-016-0213-8

Huang, Y., Jiang, L., Yang, H., Wu, L., Xu, N., Zhou, X., & Li, J. (2019). Variations of Wnt/beta-catenin pathway-related genes in susceptibility to knee osteoarthritis: A three-centre case-control study. J Cell Mol Med, 23(12), 8246–8257. doi:10.1111/jcmm.14696

Kolakshyapati, M., Flavel, R. J., Sibanda, T. Z., Schneider, D., Welch, M. C., & Ruhnke, I. (2019). Various bone parameters are positively correlated with hen body weight while range access has no beneficial effect on tibia health of free-range layers. Poult Sci, 98(12), 6241–6250. doi:10.3382/ps/pez487

Komiya, Y., & Habas, R. (2008). Wnt signal transduction pathways. Organogenesis, 4(2), 68–75. doi:10.4161/org.4.2.5851

Li, X., Lu, Y., Liu, X., Xie, X., Wang, K., & Yu, D. (2019). Identification of chicken FSHR gene promoter and the correlations between polymorphisms and egg production in Chinese native hens. Reprod Domest Anim, 54(4), 702–711. doi:10.1111/rda.13412

Lim, C. H., Lim, W., Jeong, W., Lee, J. Y., Bae, S. M., Kim, J., … Song, G. (2013). Avian WNT4 in the female reproductive tracts: potential role of oviduct development and ovarian carcinogenesis. PLoS One, 8(7), e65935. doi:10.1371/journal.pone.0065935

Niehrs, C. (2012). The complex world of WNT receptor signalling. Nat Rev Mol Cell Biol, 13(12), 767–779. doi:10.1038/nrm3470

Rufener, C., Baur, S., Stratmann, A., & Toscano, M. J. (2019). Keel bone fractures affect egg laying performance but not egg quality in laying hens housed in a commercial aviary system. Poult Sci, 98(4), 1589–1600. doi:10.3382/ps/pey544

Schwarz-Romond, T. (2012). Three decades of Wnt signalling. EMBO J, 31(12), 2664. doi:10.1038/emboj.2012.159

Stratmann, A., Frohlich, E. K., Gebhardt-Henrich, S. G., Harlander-Matauschek, A., Wurbel, H., & Toscano, M. J. (2016). Genetic selection to increase bone strength affects prevalence of keel bone damage and egg parameters in commercially housed laying hens. Poult Sci, 95(5), 975–984. doi:10.3382/ps/pew026

Tao, H., Xiong, Q., Ji, Z., Zhang, F., Liu, Y., & Chen, M. (2019). NFAT5 is Regulated by p53/miR-27a Signal Axis and Promotes Mouse Ovarian Granulosa Cells Proliferation. Int J Biol Sci, 15(2), 287–297. doi:10.7150/ijbs.29273

van Amerongen, R., Mikels, A., & Nusse, R. (2008). Alternative wnt signaling is initiated by distinct receptors. Sci Signal, 1(35), re9. doi:10.1126/scisignal.135re9

van Amerongen, R., & Nusse, R. (2009). Towards an integrated view of Wnt signaling in development. Development, 136(19), 3205–3214. doi:10.1242/dev.033910

Waghmare, I., & Page-McCaw, A. (2018). Wnt Signaling in Stem Cell Maintenance and Differentiation in the Drosophila Germarium. Genes (Basel), 9(3). doi:10.3390/genes9030127

Wang, Y., Chen, Q., Liu, Z., Guo, X., Du, Y., Yuan, Z., … Jiang, Y. (2017). Transcriptome Analysis on Single Small Yellow Follicles Reveals That Wnt4 Is Involved in Chicken Follicle Selection. Front Endocrinol (Lausanne), 8, 317. doi:10.3389/fendo.2017.00317

